# Inhibiting USP16 rescues stem cell aging and memory in an Alzheimer’s model

**DOI:** 10.1101/2020.12.21.383059

**Authors:** Felicia Reinitz, Elizabeth Chen, Benedetta Nicolis di Robilant, Bayarsaikhan Chuluun, Jane Antony, Robert C. Jones, Neha Gubbi, Sai Saroja Kolluru, Dalong Qian, Katja Piltti, Aileen Anderson, Michelle Monje, H. Craig Heller, Stephen Quake, Michael F. Clarke

## Abstract

Alzheimer’s disease (AD) is a progressive neurodegenerative disease observed with aging that represents the most common form of dementia. To date, therapies targeting end-stage disease plaques, tangles, or inflammation have limited efficacy. Therefore, we set out to identify an earlier targetable phenotype. Utilizing a mouse model of AD and human fetal cells harboring mutant amyloid precursor protein, we show cell intrinsic neural precursor cell (NPC) dysfunction precedes widespread inflammation and amyloid plaque pathology, making it the earliest defect in the evolution of disease. We demonstrate that reversing impaired NPC self-renewal *via* genetic reduction of USP16, a histone modifier and critical physiological antagonist of the Polycomb Repressor Complex 1, can prevent downstream cognitive defects and decrease astrogliosis *in vivo*. Reduction of USP16 led to decreased expression of senescence gene *Cdkn2a* and mitigated aberrant regulation of the BMP pathway, a previously unknown function of USP16. Thus, we reveal USP16 as a novel target in an AD model that can both ameliorate the NPC defect and rescue memory and learning through its regulation of both *Cdkn2a* and BMP signaling.

**Graphical Abstract:** 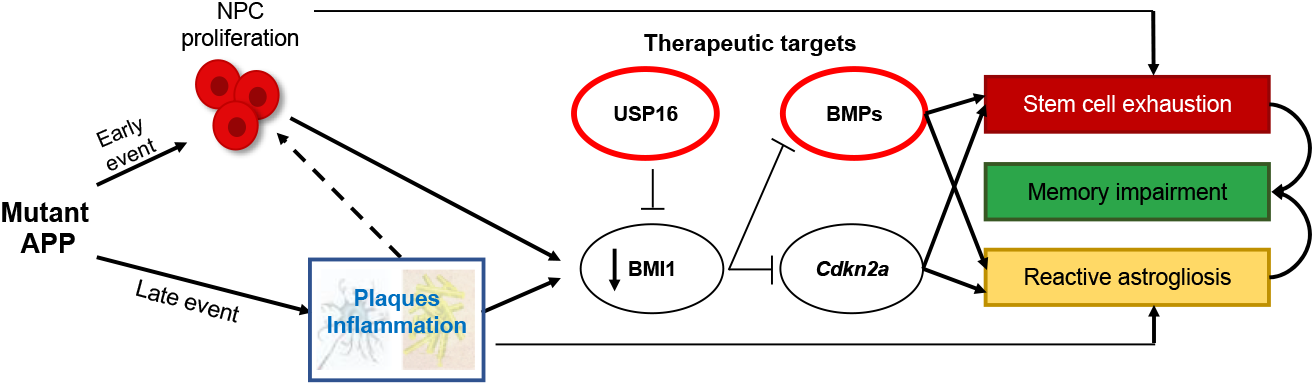

Schematic summarizing therapeutic approaches to mitigate the effects of mutant APP through targeting of *Cdkn2a*, BMI1, USP16 and BMP.

## Introduction

Alzheimer’s disease (AD) is the most common form of dementia, occurring in 10% of individuals over the age of 65 and affecting an estimated 5.5 million people in the United States (Hebert, Weuve, Scherr, & Evans, 2013). Currently there is no treatment to stop, prevent, or reverse AD (Huang & Mucke, 2012; Selkoe, 2019). Historically, AD has been understood by its end-stage disease phenotype, characterized clinically by dementia and pathologically by amyloid senile plaques and neurofibrillary tangles (Castellani, Rolston, & Smith, 2010). These traditional AD pathologies are thought to begin with amyloid plaque deposition which is associated with inflammation, increased reactive oxygen species (ROS) and neurodegeneration during aging (Akiyama et al., 2000; Glass, Saijo, Winner, Marchetto, & Gage, 2010); however, thus far, treatments to decrease formation of plaques have not significantly improved disease progression or outcomes (Selkoe, 2019).

Adult neurogenesis is thought to be compromised in AD, contributing to early dementia (Alipour et al., 2019). The decline of neural stem/progenitor cell (NPC) function in the subventricular zone (SVZ) and the hippocampus has been established in both aging (Leeman et al., 2018) and various AD mouse models (Haughey, Liu, Nath, Borchard, & Mattson, 2002; Lopez-Toledano & Shelanski, 2004; Mu & Gage, 2011; Rodriguez, Jones, & Verkhratsky, 2009; Rodriguez & Verkhratsky, 2011; Sakamoto et al., 2014; Winner, Kohl, & Gage, 2011). However, it is still not known whether these defects are cell-intrinsic resulting from changes inside the cells or extrinsic as a result of external niche factors such as inflammation. Here, we report that the neural precursor cell defects seen in an AD mouse model harboring Swedish, Dutch, and Iowa mutations in the amyloid precursor protein (Tg-SwDI) is initially cell-intrinsic and predates inflammation and widespread plaque deposition, which plays a role later in the disease. We chose the Tg-SwDI model with mutations isolated to *APP* because of early plaque deposition, cognitive deficits, and physiologic levels of mutant APP expression, rather than non-physiologic over-expressed levels as seen with other models (Davis et al., 2004). In this model, mice begin to develop amyloid deposition in brain parenchyma as early as 3 months of age and throughout the forebrain by 12 months (Davis et al., 2004; Miao et al., 2005), as well as significant cerebral amyloid angiopathy, which is thought to be a more sensitive predictor of dementia than parenchymal and amyloid plaques (Neuropathology Group. Medical Research Council Cognitive & Aging, 2001; Thal, Ghebremedhin, Orantes, & Wiestler, 2003).

Current strategies to reverse neurogenesis defects include the use of drugs (“senolytics”) that selectively remove p16^Ink4a^-positive senescent cells. Removal of the p16^Ink4a^-positive senescent cells, for instance, using a suicide gene under the regulation of the *Cdkn2a* promoter has been shown to attenuate progression of age-related decline and preserve cognitive function in both an accelerated aging AD mouse model and a tauopathy mouse model (Baker et al., 2011; Bussian et al., 2018). However, the use of a suicide gene is not directly translatable into humans, and other senolytics such as BCL2-inhibitors or the combination of Dasatinib and quercetin have toxicities which can limit their use (Amaya-Montoya, Perez-Londono, Guatibonza-Garcia, Vargas-Villanueva, & Mendivil, 2020; Zhu et al., 2015).

To circumvent these shortcomings, we targeted USP16, an upstream regulator of *Cdkn2a*, to reverse the neural precursor defect seen in the Tg-SwDI model. USP16 counteracts self-renewal regulator BMI1 by deubiquitinating histone H2A on Lysine 119 at the *Cdkn2a* locus, resulting in increased expression of protein products P16 (*Ink4a*) and P14 (*Arf*) (Adorno et al., 2013). This results in increased senescence with a concomitant decrease in self-renewal. Here, we demonstrate that inhibiting USP16 is a potential novel strategy to rescue *Cdkn2a-induced* pathologies in AD induced by both p16^Ink4a^ and p19^Arf^. To better understand its function, we probed for additional pathways regulated by *Usp16* and identified enrichment of the BMP pathway early on in the Tg-SwDI mice. The BMP pathway has been known to play a role in NPC function. Specifically, BMPR2 is a type II receptor that heterodimerizes with BMPR1a or BMPR1b and is responsible for transducing BMP signaling downstream to the SMAD proteins which translocate to the nucleus and can turn on genes related to cell fate and differentiation (Chang, Dettman, & Dizon, 2018). Levels of BMP2, 4, and 6 expression have been found to increase in the hippocampus with age (Yousef et al., 2015). Furthermore, *Bmpr2* conditional ablation in Ascl1+NSCs/NPCs or treatment with BMP inhibitor Noggin results in activation of NSCs, increased cell proliferation, and a rescue of cognitive deficits to levels comparable to young mice (Meyers et al., 2016). For the first time, we show that targeting USP16 in a mouse model of AD rescues two aberrant aging pathways, *Cdkn2a* and BMP, which can restore self-renewal of NPCs, decrease astrogliosis, and minimize cognitive decline.

### Neural precursor cell exhaustion is the earliest sign of disease in Tg-SwDI mice

Detecting disease early before fulminant pathogenesis may be crucial to developing effective diagnosis and treatment, particularly when it comes to irreversible degeneration. Therefore, we used a multimodal temporal approach including immunofluorescence staining, in vitro neurosphere assays, Luminex assays, and behavioral studies to dissect changes at the molecular, cellular, and organismal levels in both 3-4 month old and 1 year old AD mice. At 3 months of age, we found that proliferation of neural progenitor cells, marked by 5-ethynyl-2’-deoxyuridine (EdU) (Chehrehasa, Meedeniya, Dwyer, Abrahamsen, & Mackay-Sim, 2009), SOX2 and GFAP, was increased three-fold in the SVZ of Tg-SwDI mice (P=0.0153; Fig. 1A). In many tissues including the blood, pancreas, intestine and mammary gland, hyperproliferation has been linked to a premature decline in stem cell function associated with aging (Essers et al., 2009; Krishnamurthy et al., 2006; Scheeren et al., 2014). Using extreme limiting dilution analysis (ELDA) of neurosphere-formation from single cells (Hu & Smyth, 2009; Pastrana, Silva-Vargas, & Doetsch, 2011) we discovered that 3-4 month old Tg-SwDI mice had significantly less regenerative potential of the SVZ cells than that of healthy age-matched control mice (neurosphere-initiating cell (NIC) frequencies: 1 in 14.5 vs 1 in 7.5, respectively, P=0.00166, Fig. 1B).

**Figure 1:**
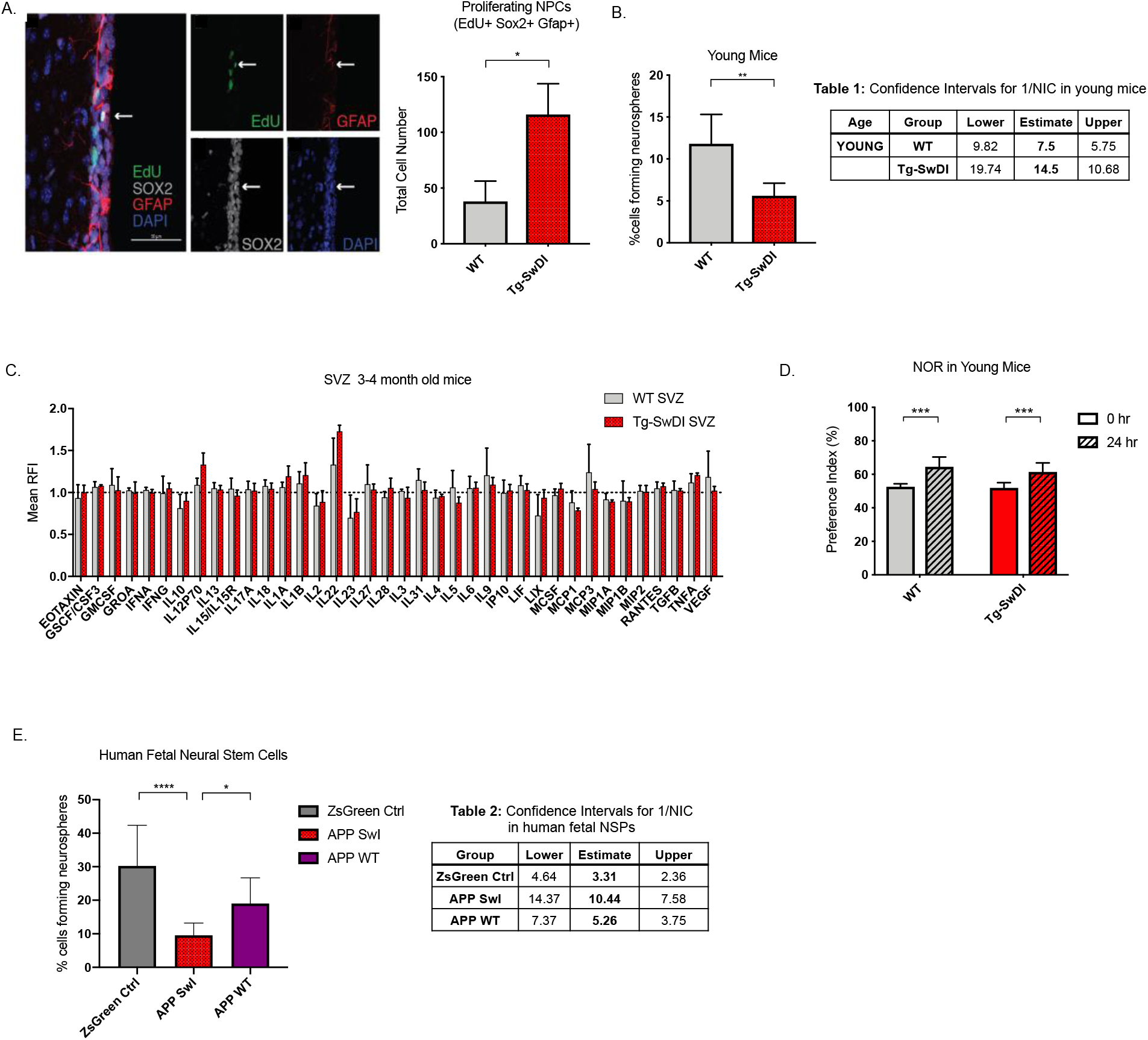
Defects in self-renewal and hyperproliferation in Tg-SwDI mice predate cognitive deficits and widespread inflammation. **(A)** Representative 40X confocal images of the SVZ stained for EdU, GFAP and SOX2. 3-4 month old mice underwent intraperitoneal injections every day for 6 days with EdU and the analysis was performed four weeks afterward to capture all true activated daughter stem cells that would maintain the niche without further differentiation or migration. Count of proliferating NPCs, as cells positive for EdU, GFAP, SOX2 and DAPI, is shown in the panel on the right (n=3 mice). Data are presented as mean ± SEM. **(B)** Limiting Dilution assays were performed using a series of dilutions (1 cell/well, 5 cells/well, 25 cells/well, 125 cells/well) derived from neurospheres from the SVZ of 3-4 month old mice (n=3). The graph shows the percentage of neurosphere-initiating cells (NIC) ± upper and lower estimates converted values calculated by ELDA. Twelve technical replicates of each dilution were counted. **Table 1:** summarizes the upper and lower confidence intervals and estimates of 1/NIC for the different genotypes calculated by ELDA. **(C)** Cytokine levels measured by Luminex array from the SVZ of young 3-4 months mice. No differences have been observed at this age (n=3 mice for each genotype). See also Fig. S1. Data are presented as mean ± SD. **(D)** NOR 24-hour testing in mice at 3 months of age showed no signs of cognitive impairment in the Tg-SwDI mice with a preference index comparable to that of WT indicating both genotypes had intact object discrimination (P=0.001 for WT and P=0.0099 for Tg-SwDI, n=7-10 mice in each group). Data are presented as mean ± SEM. **(E)** Left panel shows percentage of NIC ± upper and lower estimates for human fetal neurospheres infected with pHIV-Zsgreen, wild type APP (APP WT), or APP SwI (Swedish and Indiana mutations). Table 2 (right): Lists the estimated stem cell frequencies and ranges for each group, calculated using the ELDA software (n=3 separate infections and limiting dilution experiments with 12 technical replicates for each limiting dilution of 1 cell/well, 5 cells/well, 25 cells/well, 125 cells/well) (****P=3.7e^−6^, *P=0.00507).

A well-established prominent phenotype associated with AD is inflammation, although it is not clear when brain inflammation can first be detected. To explore whether an inflammatory signature was present by 3-4 months of age, we employed a Luminex screen to assess the presence of an array of cytokines and other inflammatory markers. We looked at the SVZ, hippocampal dentate gyrus (DG), and cortex in 3-4 month old mice, but found no significant differences in inflammatory markers between Tg-SwDI and wild type mice in any of these regions (Fig. 1C and Fig. S1). To explore further, we measured mRNA levels of *Ptgs2* (COX2), *Tnfα, Il6* and *Illβ* utilizing qPCR, but found no significant differences between WT and Tg-SwDI in either the SVZ, DG or Cortex (Fig S1C).

One of the most debilitating features of AD is memory impairment and progressively diminished cognitive function. Although Tg-SwDI mice are known to exhibit these features, there was no evidence of cognitive impairment in 3-4 month old mice when subjected to novel object recognition (NOR) training and subsequent testing after 24 hours (Ennaceur & Delacour, 1988) (Fig. 1D).

Given the prominent early aberrant self-renewal phenotype in the young Tg-SwDI mouse model, we investigated whether or not expression of mutant *APP* in human NPCs might also cause a self-renewal defect. We therefore infected human fetal neurospheres with a lentiviral construct for either pHIV-Zsgreen alone, pHIV-Zsgreen with wild type *APP*, or pHIV-Zsgreen with Swedish and Indiana *APP* mutations (APP SwI). Employing the same limiting dilution assay as before, we found diminished NIC frequency of mutant *APP*-infected human neurospheres compared to cells infected with the empty vector or with wild-type *APP* (1 in 10.44 vs 1 in 3.31 and 1 in 10.44 vs 1 in 5.26, respectively, P=3.7e^−06^ and P=0.00507; Fig. 1E and Table 2). This result suggests that the self-renewal defect seen in the Tg-SwDI Alzheimer’s model is cell-intrinsic and not specific to the Swedish, Dutch and Iowa mutations, but also more broadly seen with other *APP* mutations. Importantly, these results also demonstrate that the effects observed in NPCs derived from a genetic mouse model can be robustly recapitulated in human NPCs expressing mutant *APP*.

### Modest aging in Tg-SwDI accelerates NPC exhaustion and astrogliosis prior to inflammation

To explore progression of the disease with aging, we next looked at what phenotypic changes occurred in 1 year old Tg-SwDI mice, including self-renewal, astrogliosis and inflammation. The defect in self-renewal that was observed in the 3-4 month old Tg-SwDI mice was exacerbated in the 1 year old Tg-SwDI mice (1 in 35.5 for Tg-SwDI mice vs 1 in 17.6 for WT mice, P=0.00625; Fig. 2A and Table 3). The NIC capacity in 1 year old controls was similar to that of young Tg-SwDI mice (1 in 17.6 vs 1 in 14.5, respectively, compare Tables 1 to Table 3), emulating an accelerated aging phenotype with mutant APP. Although we observed continuous passaging of neurospheres diminished with age alone in healthy control mice (P=0.021; Fig. S2A), aging more severely impacted the Tg-SwDI mice. In line with these findings, expression of the well-studied gene, *Cdkn2a*, known for its increased expression with aging and critical function of inhibiting stem cell self-renewal during development and throughout the lifespan, was increased with aging and even more so in the Tg-SwDI cortex (Fig. 2B).

**Figure 2:**
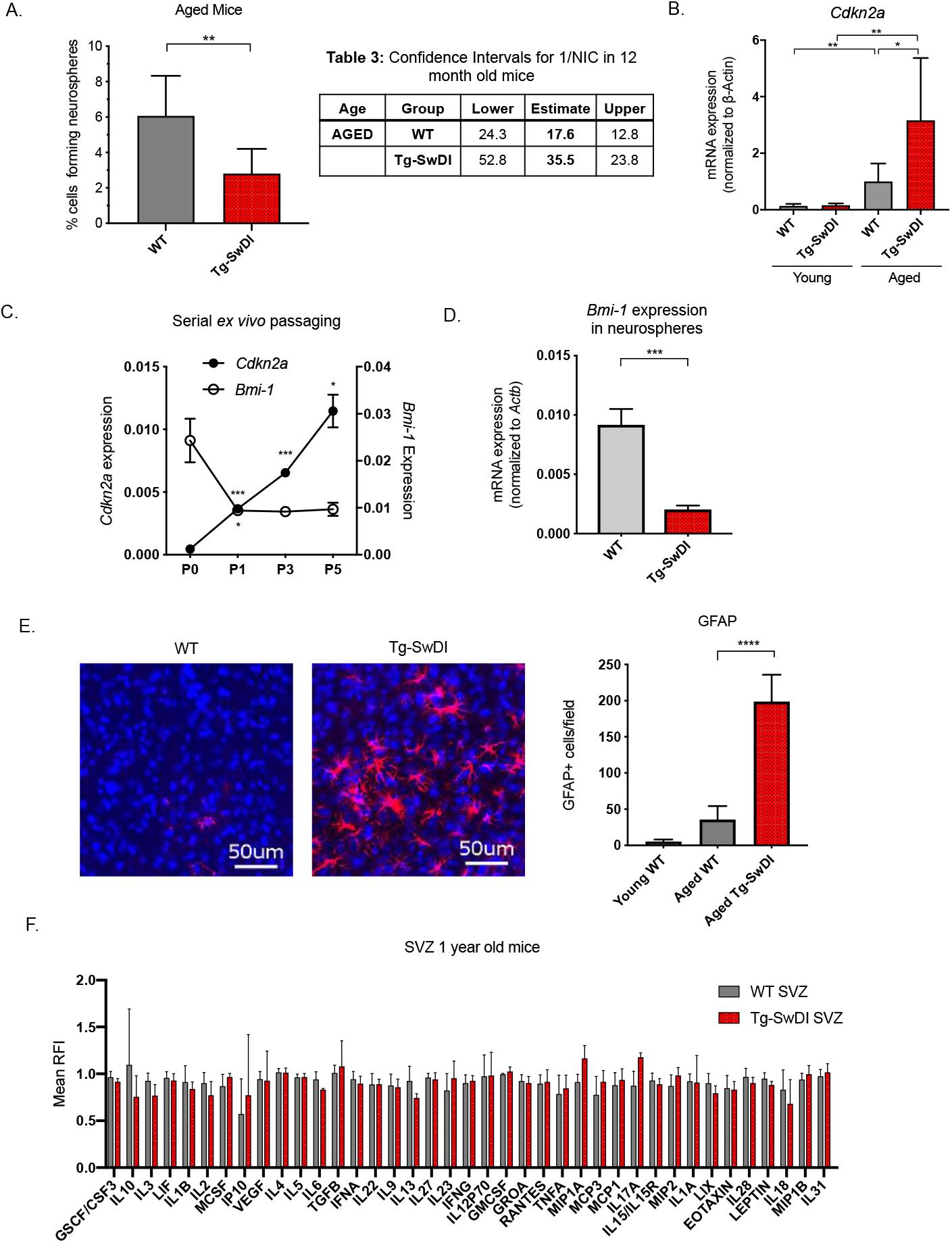
Accelerated aging phenotype seen in Tg-SwDI mice with exacerbated self-renewal and astrogliosis. **(A)** Limiting Dilution assays were performed using single cells derived from the SVZ of 1 year old mice. The graph shows the percentage of neurosphere-initiating cells calculated by ELDA. Table summarizes the estimates of 1/NIC for the different genotypes calculated by ELDA. **(B)** mRNA levels of *Cdkn2a* in the cerebral cortex of young (3-4 month old) and aged (12 month old) mice were measured by RT-qPCR. Ct values were normalized to β-Actin. (WT = wild-type littermate; *Cdkn2a:* n=6-7 mice each genotype). A one-way ANOVA showed significant differences between the groups (P=0.0063 between Aged WT and Aged Tg-SwDI). Data are presented as mean ± SD. **(C)** mRNA levels of *Cdkn2a* and *Bmi1* during WT neurosphere serial passaging in vitro. Every 4-8 days (depending on neurosphere size), neurospheres were dissociated and re-plated at a density of 10 cells/uL. (n=3 mice, mice were aged 3-4 months). Data are presented as mean ± SD. **(D)** *Bmi1* expression levels measured by RT-qPCR in neurospheres derived from the SVZ of WT or Tg-SwDI mice at 3^rd^ passage (mice aged 3-4 months). Data are presented as mean ± SD. **(E)** Anterior sections from 9-12 month old mice were stained for GFAP+ cells in the cortex. Four regions per section and three sections per mouse were counted (n=4 mice each group). A one-way ANOVA showed significant differences between the groups (P<0.0001 between aged WT and Tg-SwDI). Data are presented as mean ± SD. **(F)** Cytokine levels measured by Luminex array from the SVZ of 1 year old mice. No differences were observed at this age. (n = 3 mice per genotype). See also Fig. S2B. Data are presented as mean ± SD.

Reactive astrogliosis, the abnormal increase and activation of astrocytes that can drive degeneration of neurons, has also been linked to both AD disease pathogenesis (Osborn, Kamphuis, Wadman, & Hol, 2016) and to the *BMI1/Cdkn2a* pathway. Specifically, Zencak and colleagues showed increased astrogliosis in the brain of *Bmi1^-/-^* mice (Zencak et al., 2005). As a secondary effect of the reduced passaging capacity of neurospheres that accompanies aging, we observed an increase in *Cdkn2a* expression in neurospheres along with a decrease in *Bmi1* expression (Fig. 2C). In neurospheres derived from the SVZ of Tg-SwDI mice, we observed an even greater decrease in *Bmi1* expression (Fig. 2D). Furthermore, we observed that age-related astrogliosis, marked by GFAP+ cells, significantly increased in Tg-SwDI mice (P = 0.0010, respectively; Fig. 2E).

Often associated with astrogliosis is neuroinflammation (Frost & Li, 2017). As previously tested in the 3-4 month old mice, we looked for the presence of inflammatory cytokines using the Luminex array in 1 year old mice. We hypothesized that inflammation might explain some of the aging phenotypes observed thus far. However, even at 1 year old, there were no significant inflammatory differences in the SVZ, DG, or cortex between the WT and Tg-SwDI mice (Fig. 2F, Fig. S1C and S2B). Taken together with our data in human NPCs expressing mutant *APP*, the phenotypes seen at both 3-4 months old and 1 year old suggest a cell intrinsic defect rather than an outcome of cell extrinsic factors like inflammation. Moreover, through these data illustrating the changes that occur with aging in Tg-SwDI mice, we identified an NPC defect as the earliest indication of disease.

### Self-renewal defects are rescued by Usp16 and Cdkn2a modulation

Neural precursor cells function through a number of genetic and epigenetic components, and one of the well described master regulators is *Cdkn2a*, a gene tightly regulated by BMI1 (Bruggeman et al., 2005). When we crossed the Tg-SwDI mouse with a *Cdkn2a* knockout mouse (Tg-SwDI/*Cdkn2a*^-/-^) and performed limiting dilution assays in SVZ cells from 3-month old mice, a complete restoration of the NIC frequency in the Tg-SwDI/*Cdkn2a*^-/-^ cells compared to age-matched Tg-SwDI cells was observed (P=7.7e^−05^; Fig. 3A). This NIC rescue was also observed in hippocampal cells cultured from microdissection of the dentate gyrus (DG) (P=2.09e^−9^, Fig. 3B). These results demonstrate that impairment of NPC regeneration, as measured by NIC frequencies, is a function of aging that is accelerated by *APP* mutations and is mediated through *Cdkn2a*, a known regulator of NPC self-renewal (Molofsky et al., 2003).

**Figure 3:**
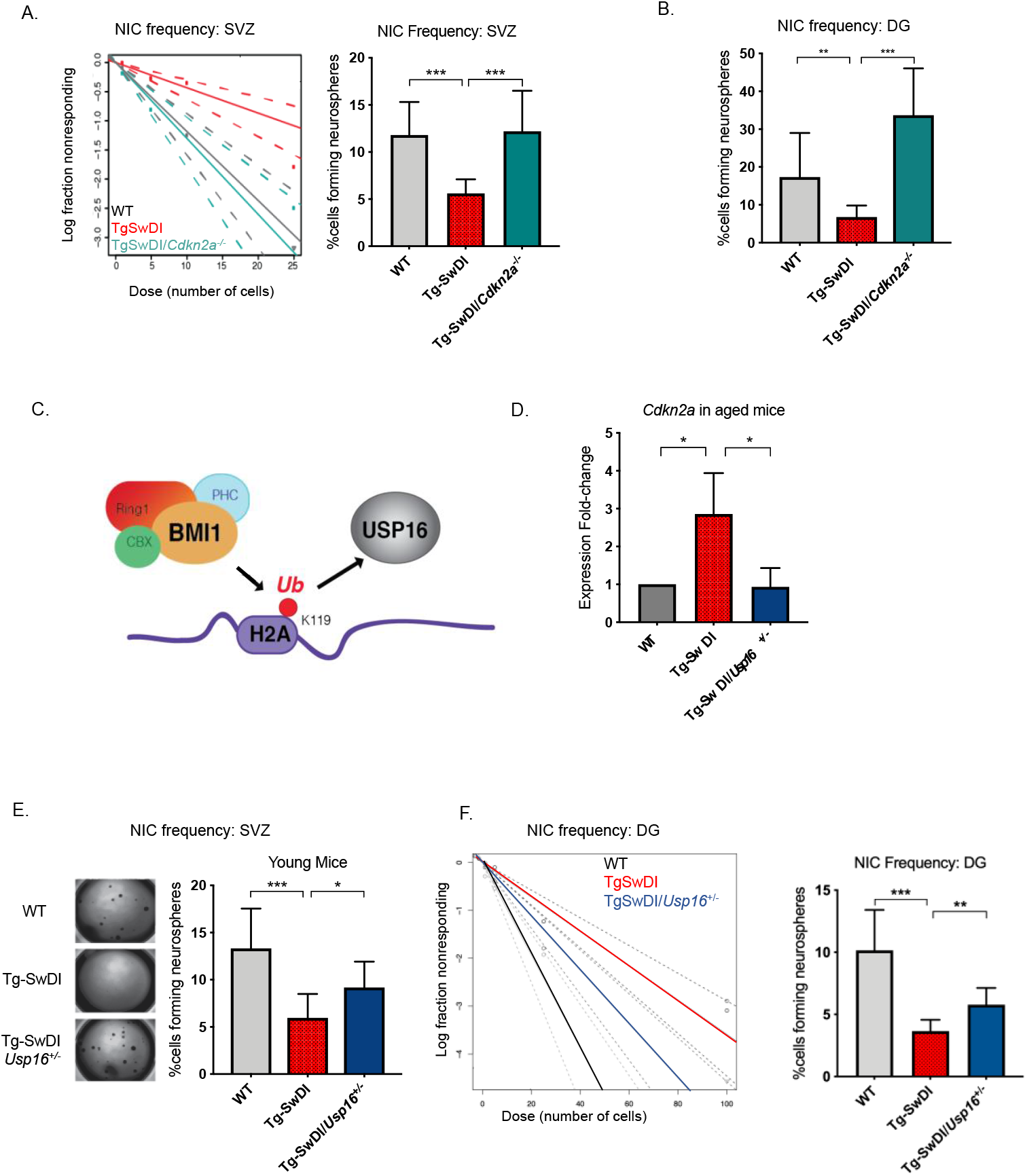
Usp16 haploinsufficiency normalizes *Cdkn2a* expression and restores self-renewal in Tg-SwDI NPCs. **(A)** Left panel shows ELDA graph of limiting dilution assay comparing SVZ from young WT, Tg-SwDI and Tg-SwDI/*Cdkn2a^-/-^* mice. The bar graph shows the NIC frequencies in SVZ and in the dentate gyrus **(B)** as percentages of total cells with error bars indicating the upper and lower values. Mice were 3 months old when sacrificed; experiment done after 3^rd^ passage of NSPs. **(C)** Schematic summarizing the role of *Bmi1* in ubiquitinating histone H2A at the *Cdkn2a* locus and the role of USP16 as its natural antagonist, suggesting that USP16 inhibition could have an effect on self-renewal. **(D)** RT-qPCR of Cdkn2a in the cerebral cortex of old Tg-SwDI mice shows mRNA levels were rescued by *Usp16* haploinsufficiency (n=3). Ct-values were normalized to β-actin. A one-way ANOVA showed significant differences between the groups (P=0.0365 between WT and Tg-SwDI and P=0.0318 between Tg-SwDI and Tg-SwDI/Usp16^+/-^). Data are presented as mean ± SD. **(E)** Left panel shows 1X representative photographs of neurospheres grown in 96-well dish after 2 weeks of culture. The bar graph shows the NIC frequencies as percentages of total cells comparing WT, Tg-SwDI and Tg-SwDI/Usp16^+/-^ mice. Mice were 3 months old. **(F)** ELDA graph of limiting dilution assay comparing hippocampal cells obtained from the dentate gyrus on left. Bar graph on right shows percent of cells forming neurospheres (**P = 0.00233 and ***P = 9.9e^−9^).

Unfortunately, mutations or loss of function in the *Cdkn2a* gene eventually leads to tumor formation, making it not feasible to perform limiting dilution experiments in aged *Cdk2na* knockout mice and also making it less than ideal to target therapeutically (Hussussian et al., 1994). Upstream of *Cdkn2a* is USP16, an antagonist of BMI1 and a de-repressor of *Cdkn2a* that acts through the enzymatic removal of ubiquitin from histone H2A (Fig. 3C) (Adorno et al., 2013; Joo et al., 2007). We predicted that downregulation of *Usp16* would increase BMI1 function to counteract the effects of mutant APP similar to what we observed with knockout of *Cdkn2a*. This is supported by previous data that showed overexpression of *USP16* in human-derived neurospheres led to a marked decrease in the formation of secondary neurospheres (Adorno et al., 2013). To test this, we crossed Tg-SwDI with *Usp16^+/-^* mice to generate Tg-SwDI/*Usp16^+/-^* mice, which do not show tumor formation. We found that Tg-SwDI mice express greater than two-fold more cortical *Cdkn2a* than both WT and Tg-SwDI/*Usp16^+/-^* mice, for which expression levels were very similar (P=0.0365 and P=0.0318, respectively, Fig. 3D). Limiting dilution experiments of cells isolated from the SVZ and DG of the hippocampus showed that Tg-SwDI/*Usp16^+/-^* mice had significantly greater NIC frequencies, partially rescuing the self-renewal defect seen with mutant APP (P=0.0492 and P=0.00233, respectively; Fig. 3E and F). Similar to the NIC rescue in the Tg-SwDI/*Cdkn2a^-/-^*, these data show cell-intrinsic impaired self-renewal in the Tg-SwDI model of familial AD, and that reversal of this impairment is possible through targeting *Cdkn2a* upstream regulator, USP16.

### RNA-seq data reveals enriched BMP signaling in Tg-SwDI mice that is rescued by Usp16 haploinsufficiency

To delineate potential self-renewal pathways that might contribute to the defect and rescue of Tg-SwDI NPCs and Tg-SwDI/*Usp16^+/-^* NPCs, respectively, we performed single-cell RNA-seq and gene set enrichment analysis (GSEA) on lineage depleted primary FACS-sorted CD31^−^CD45^−^ Ter119^−^CD24^−^ NPCs from Tg-SwDI, WT, and Tg-SwDI/*Usp16^+/-^* mice at 3-4 months and 1 year of age (Fig. 4A) (Mootha et al., 2003; Subramanian et al., 2005). Using the GSEA Hallmark gene sets, we found only three gene sets that were enriched in Tg-SwDI mice over WT mice and rescued in the Tg-SwDI/*Usp16^+/-^* mice at both ages: TGF-ß pathway, oxidative phosphorylation, and Myc Targets (Table 4). The TGF-ß pathway consistently had the highest normalized enrichment score in pairwise comparisons between Tg-SwDI vs WT and Tg-SwDI vs Tg-SwDI/*Usp16^+/-^* of the three rescued pathways (Table 5). In looking specifically at the leading-edge genes contributing to the enrichment plots of the TGF-ß pathway, we found upregulation of BMP receptors and *Id* genes which are known to be involved in the BMP pathway, a sub-pathway of the greater TGF-ß pathway (Fig. 4B). Heatmaps of average normalized single-cell gene expression showed BMP receptors as the highest expressed TGF-ß receptors in the sorted cells, with genes such as *Bmpr2, Bmpr1a, Id2*, and *Id3* upregulated in Tg-SwDI mice and rescued in Tg-SwDI/*Usp16^+/-^* mice (Fig. 4C). These data suggest that USP16 may regulate neural precursor cell function in part through the BMP pathway.

**Figure 4:**
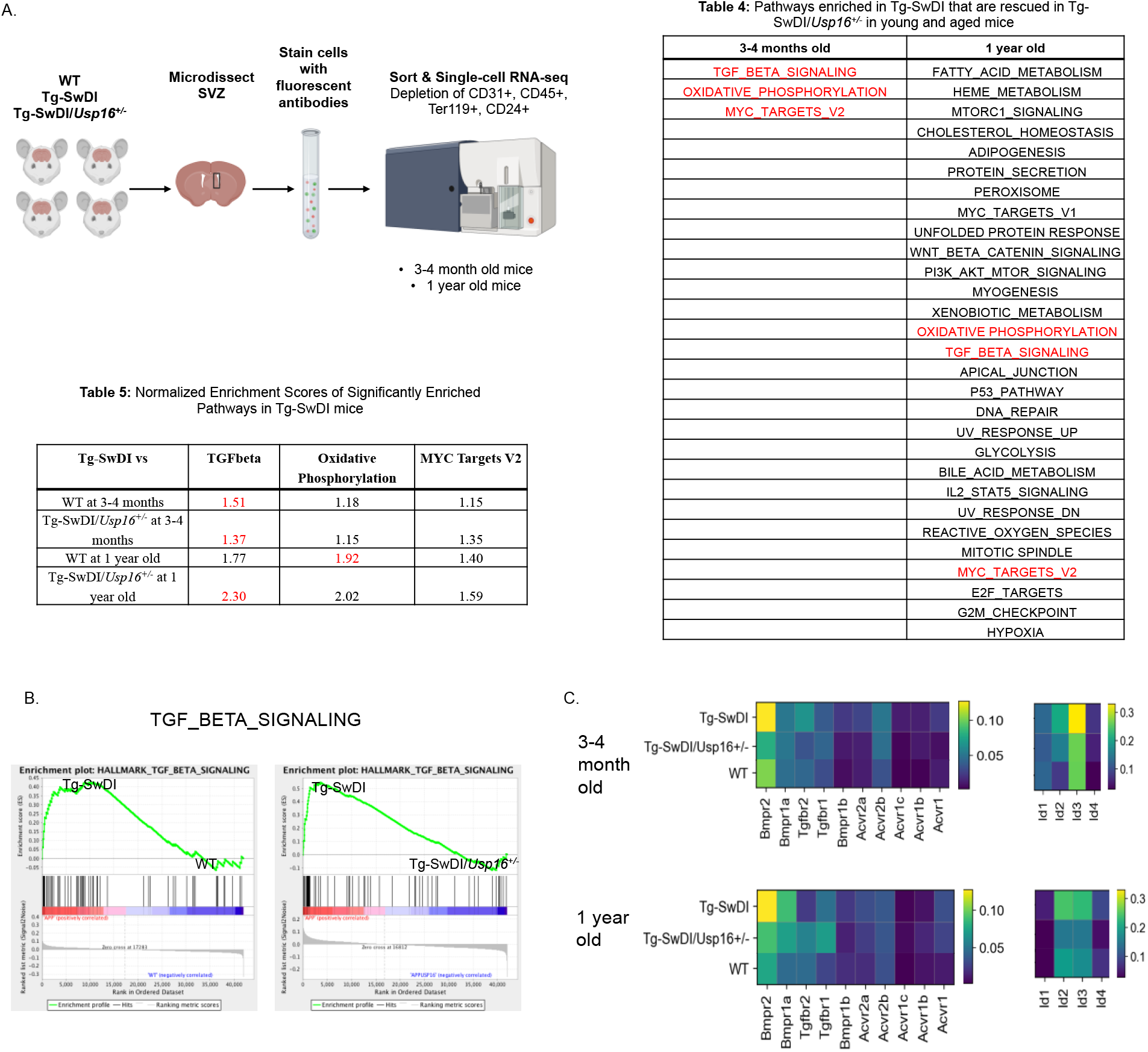
BMP signaling is enriched in Tg-SwDI and decreases with *Usp16* haploinsufficiency. **(A)** Lineage^−^CD24^−^ NPCs were FACS-sorted from the SVZ of 4 mice each of the different genotypes and processed for single-cell RNA-sequencing. **Table 4:** GSEA analysis from single-cell RNA-seq data shows pathways enriched in Tg-SwDI mice compared to WT and rescued in Tg-SwDI/ *Usp16^+/-^* mice, ordered top to bottom from smallest FDR q-val (most significant) to largest FDR q-val (least significant). (n=4 for each genotype at each time point; FDR<25%). Pathways in common to both age groups are in red. **Table 5:** Normalized enrichment scores of significantly enriched pathways in Tg-SwDI mice compared to WT or Tg-SwDI/*Usp16*^+/-^ mice at different time points. TGF-ß, Oxidative phosphorylation, and MYC Targets V2 were selected as they were rescued in both 3-4 months and 1 year old mice by Usp16 haploinsufficiency. Highest normalized enrichment scores of each comparison are in red text. **(B)** Enrichment plots show TGF-ß signaling pathway as enriched in Tg-SwDI mice and rescued by *Usp16* haploinsufficiency. Normalized Enrichment Score (NES) for left panel is 1.77 with FDR-q value = 0.008; NES for right panel is 2.30 with FDR q-value < 0.001. **(C)** Heatmaps showing averaged normalized single-cell gene expression of elements of the TGF-ß pathway; elements of the BMP pathway, a sub-pathway of the TGF-ß pathway, are specifically enriched in Tg-SwDI mice.

### BMPR inhibition rescues stem cell defects and abolishes increased phospho-SMAD 1/5/8

To confirm the functional significance of the BMP pathway in APP-mediated self-renewal defects, we measured the effects of modulating BMP pathway activity *in vitro* in human fetal NPCs expressing *APP* with Swedish and Indiana mutations (APPSwI). First, we measured levels of phosphorylated-SMAD (pSMAD) 1, 5, and 8, known readouts of BMP activity, and found they were significantly increased in the mutant neurospheres compared to control (P=0.0001, Fig. 5A). Treatment of the neurospheres with the BMP receptor inhibitor LDN-193189, a specific inhibitor of BMP-mediated SMAD1, SMAD5, and SMAD8 activation, substantially decreased pSMAD 1/5/8 in APPSwI NPCs (P<0.0001, Fig. 5B and C) (Yu et al., 2008). Furthermore, when we treated neurospheres expressing mutant *APP* with LDN-193189 for a week, the number of colonies originating from those cells were similar to control cells and significantly higher than untreated mutant *APP* neurospheres (Fig. 5D and E). Notably, LDN-193189 had minimal impact on Zsgreen control neurosphere growth (Fig. 5E). This finding demonstrates that the decrease in NIC frequency observed with mutant APP could be explained by the upregulation of BMP signaling. Moreover, BMPR inhibition rescues this defect in cells overexpressing mutant *APP* at doses that have no toxic effect on healthy cells. Altogether, these data reveal that BMP signaling enrichment is recapitulated in human NPCs expressing mutant *APP*, and that BMP inhibition normalizes the stem cell defect.

**Figure 5.**
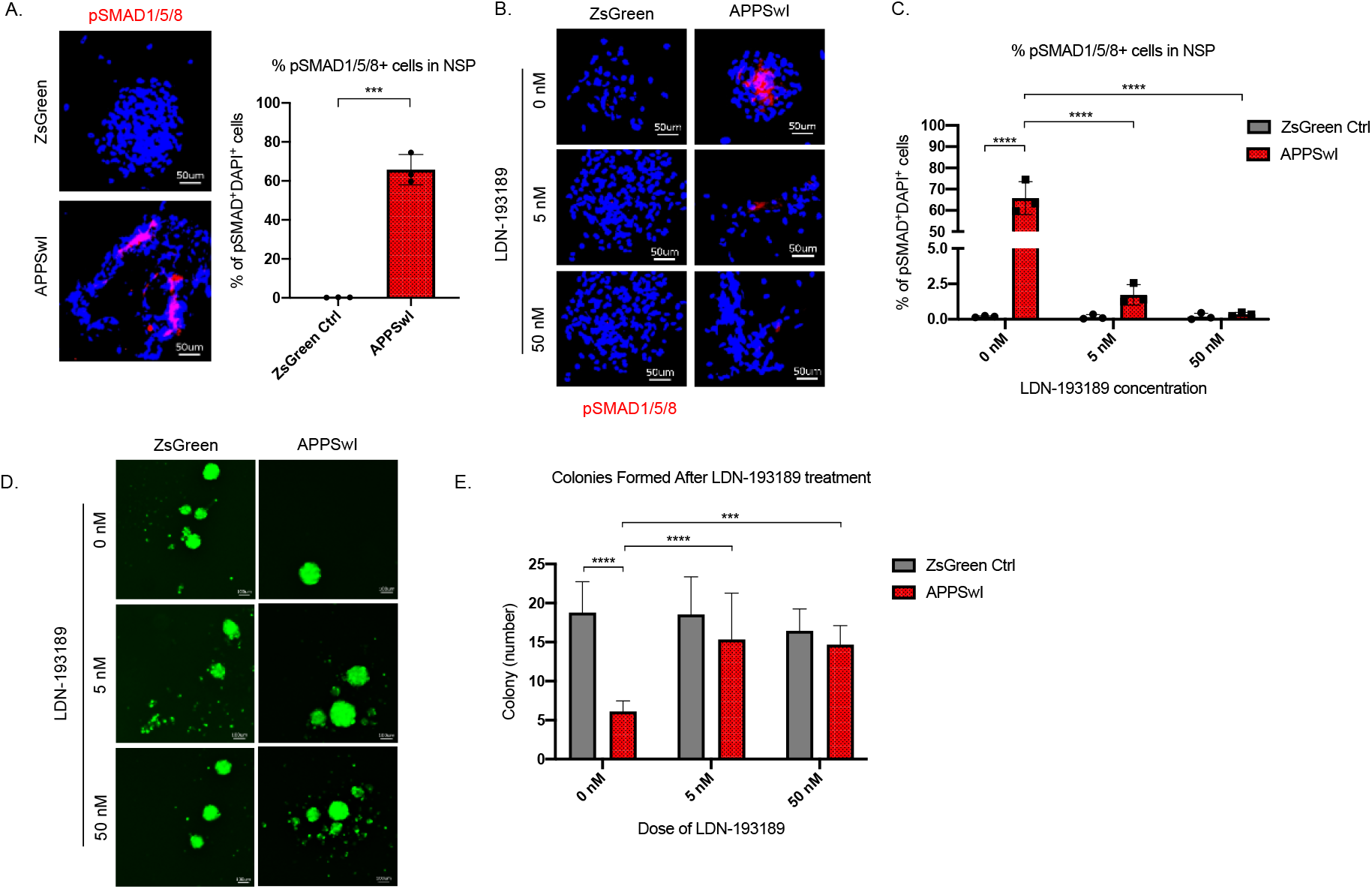
BMPR inhibition rescues mutant APP mediated self-renewal defects in human neurospheres. **(A)** Left panel shows representative images of phospho-Smad 1/5/8 staining in mutant APP-infected human fetal neurospheres compared to Zsgreen controls. Scalebar is 50 μm. Right panel shows quantification of DAPI and phospho-Smad1/5/8 co-stained cells in each group. Data are presented as mean ± SD. **(B)** Representative images of phospho-Smad1/5/8 staining in neurospheres treated with LDN-193189 for 1 week. Scalebar is 50 μm. **(C)** Quantification of phospho-SMAD 1/5/8 after treatment with different doses of LDN-193189. A two-way ANOVA revealed significant differences between the groups (**** for P<0.0001). Data are presented as mean ± SD. **(D)** Representative images of in vitro colonies of mutant APP- and Zsgreen-infected human fetal neurospheres after 1 week of LDN-193189 treatment. Scalebar is 100 μm. **(E)** Quantification of the colonies in (D). A two-way ANOVA revealed significant differences between groups (**** for P<0.0001 and *** for P=0.0003). Data are presented as mean ± SD.

### Astrogliosis is reduced and cognitive function is restored in Tg-SwDI/Usp16^+/-^ mice

Having identified USP16 as a target to modulate two critical pathways affected by mutations in APP, *Cdkn2a and BMP*, we investigated further Usp16’s potential effects on downstream pathophysiological markers of AD that are recapitulated in the Tg-SwDI model such as astrogliosis, inflammation, amyloid plaques and memory. Astrogliosis, marked by GFAP+ cells, was increased throughout the cortex in Tg-SwDI mice and was significantly reduced with *Usp16* haploinsufficiency (Fig. 6A).

**Figure 6:**
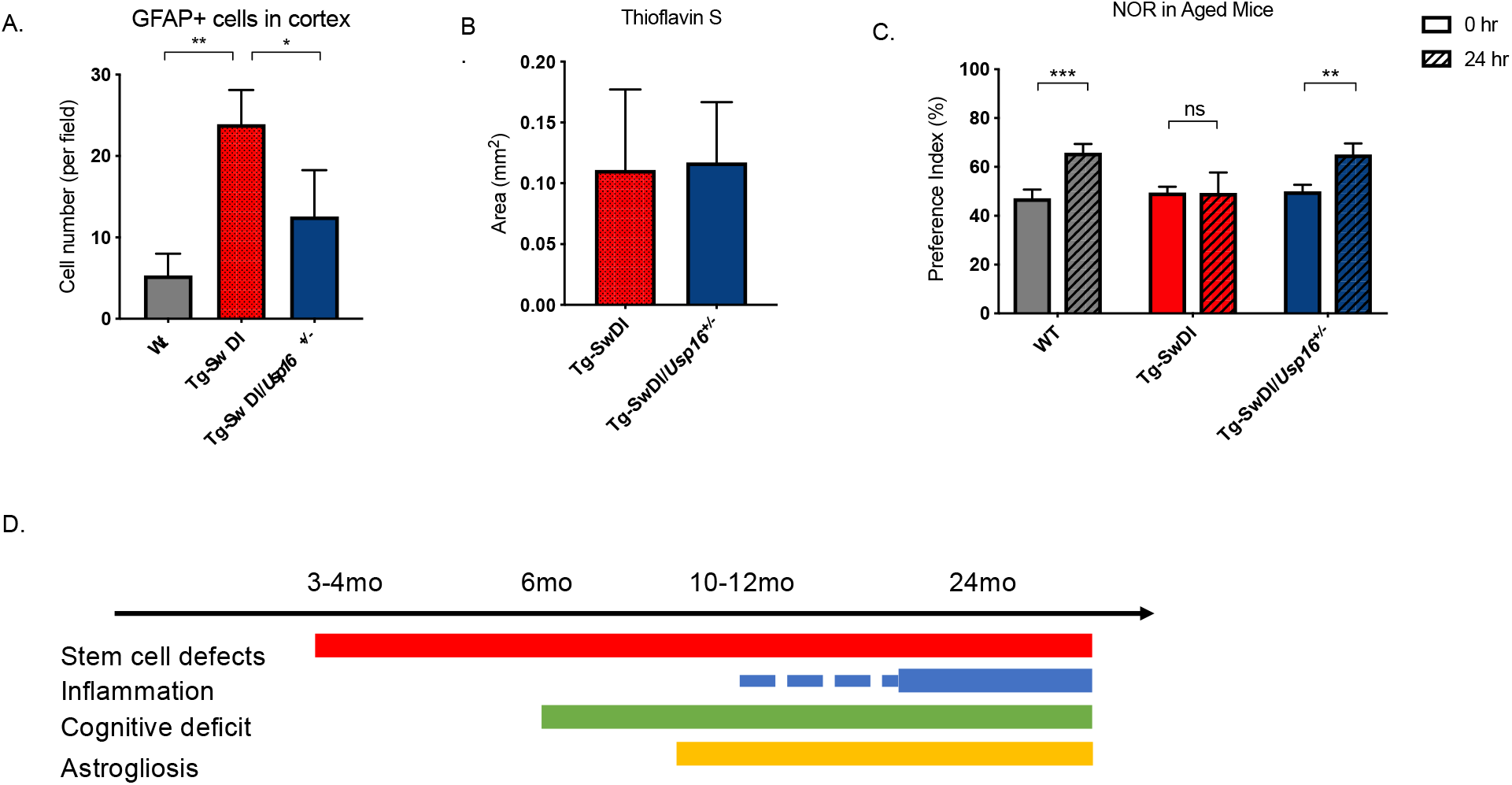
Astrogliosis, cognitive deficits, but not amyloid plaque burden are some of the processes rescued in Tg-SwDI/*Usp16^+/-^* mice. **(A) Graph shows total number of GFAP+ cells (**anterior cortical sections were stained and analyzed from 9-12 month old mice. Four different images per sections and three sections per mouse were counted; n=4). A one-way ANOVA showed significant differences between the groups (P=0.0012 between WT and Tg-SwDI and P=0.0188 between Tg-SwDI and Tg-SwDI/*Usp16^+/-^*). Data are presented as mean ± SD. **(B)** Quantification of area covered by plaques using thioflavin S staining in Tg-SwDI and Tg-SwDI/*Usp16^+/-^* mice shows no difference between the two genotypes (10-month-old mice). Data are presented as mean ± SEM. **(C)** NOR 24-hour testing in mice at 6 months of age showed the earliest signs of cognitive impairment in the Tg-SwDI mice with a preference index of 49%, while WT and Tg-SwDI/Usp16^+/-^ mice had preference indexes >65% indicating intact object discrimination (P = 0.001 for WT and P = 0.0099 for Tg-SwDI/Usp16^+/-^, n = 7-10 mice per genotype). Data are presented as mean ± SEM. **(D)** Schematic summarizing the temporal effects of mutant APP demonstrated in this manuscript.

Amyloid plaques are one of the defining features of AD, and controversy exists concerning the effect of plaques on cognitive decline. Mutations in *APP* lead to amyloid plaque deposition throughout the brain as seen in the aged Tg-SwDI mice. However, no difference was observed in plaque burden, demonstrated by Thioflavin S staining, in the age-matched Tg-SwDI/*Usp16^+/-^* mice (Fig. 6B). In addition, a Luminex screen of the Tg-SwDI/*Usp16^+/-^* mice also did not reveal significant differences in the levels of inflammatory cytokines from any of the groups (Fig. S3).

As expected, when studying the cognitive decline in the Tg-SwDI cohort, we found that the Tg-SwDI cohort exhibited impaired performance in the NOR task as early as 6 months of age, with preference indexes (P.I.s) that were not significantly different 24 hours after training, indicating no memory of the familiar object (Fig. 6C). The Tg-SwDI/*Usp16^+/-^* mice performed equally to their age-matched wild-type controls indicating memory of the familiar object with P.I.s in the 65-70% range (P=0.0099; Fig. 6C). Long-term memory impairment in Tg-SwDI mice and rescue in Tg-SwDI/Usp16+/-mice was further supported by the Barnes maze (BM) where Tg-SwDI mice spent more time exploring off-target quadrants and Tg-SwDI/Usp16+/-mice spent more time in the target quadrant (P = 0.0128 and P = 0.0251, respectively; Fig. S4). These data indicate that although modulating *Usp16* gene dosage does not affect amyloid plaque burden, it ameliorates stem cell self-renewal defects which may be the earliest indication of pathology, as well as reactive astrogliosis and some of the cognitive defects in these mice that occur later (Fig. 6D).

## Discussion

Numerous studies have sought to target processes such as inflammation, amyloid plaque accumulation, and ROS to AD pathologies in both humans and mouse models (Gjoneska et al., 2015). The lack of efficacy in trials that utilize therapies directed against amyloid and inflammatory pathways even when initiated early in the disease (Imbimbo, Solfrizzi, & Panza, 2010) suggests that other mechanisms are at play. If so, identification of these other disease mechanisms is needed to develop effective treatments (Aisen, 2008; Doody et al., 2013; Green et al., 2009; Group et al., 2008; Salloway et al., 2014).

One of the primary findings in this study is that an NPC defect predates the development of inflammation in a mutant APP model, and that this defect is cell intrinsic. We further show this cell intrinsic NPC defect is reproduced in human fetal NPCs expressing *APP* Swedish and Indiana mutations. This suggests that our findings are more broadly translatable to other *APP* mutations seen in familial human Alzheimer’s Disease.

The NPC defect that we discovered is partly regulated by *Cdkn2a*, a central component of aging, decreased neurogenesis, and differentiation of NPCs (Abdouh et al., 2012; Molofsky et al., 2006). As inhibiting *Cdkn2a* can result in tumor formation, we explored modulation of its upstream regulator, USP16. When we inhibited USP16 by making Tg-SwDI mice haploinsufficient for *Usp16* (Tg-SwDI/*Usp16^+/-^*), we found a rescue in the self-renewal of NPCs as early as 3 months of age.

We also demonstrated a new role for USP16 in regulating the BMP pathway, a mechanism independent of *Cdkn2a*. Previously, Gargiulo *et al*. found that self-renewal gene *Bmi1*, whose PRC1 activity is counterbalanced by USP16, represses BMP signaling (Gargiulo et al., 2013). NPCs from *Bmi1* knockout mice treated with BMP4 experience even further growth arrest than those untreated (Gargiulo et al., 2013). Furthermore, Kwak, Lohuizen and colleagues showed that treatment of human neural stem cells with secreted APPα or overexpression of *APP* promoted phosphorylation of SMAD 1/5/8 and induced massive glial differentiation (by expression of GFAP) through the BMP pathway (Kwak, Hendrix, & Sugaya, 2014). In line with this, our results reveal expression of mutant APP in human fetal NPCs induced phosphorylation of SMAD 1/5/8 and reduced neurosphere colony formation that was rescued by a BMP receptor inhibitor. Importantly, our data extends their findings of astrogliosis to an *in vivo* mouse model of Alzheimer’s disease. Interestingly, BMI1 regulates both *Cdkn2a* and BMPs independently, and BMI1 expression was shown to be decreased in AD patients compared to age-matched controls (Flamier et al., 2018).

Understanding the pathophysiology of a disease is critical to developing therapeutic targets and designing intervening therapies. Here, we present USP16 as a potential therapeutic target acting on both BMP and *Cdkn2a* pathways independently (summarized in the summary schematic). It is important to note that USP16 reduction also reduced astrogliosis and restored cognitive function as measured by the NOR test, independently of plaques and inflammation. The increase in astrogliosis and impaired cognitive function seen in this AD model are purely attributable to mutant APP as Tg-SwDI mice do not develop neurofibrillary tangles that require mutations in tau (Wilcock et al., 2008). Thus, therapeutic strategies that combine targeting USP16, which effectively rescues the mutant APP-induced cell intrinsic damage, with agents that target extracellular plaque formation, neurofibrillary tangles and/or inflammation may improve treatments for AD.

## Acknowledgments

We thank several individuals including Siddhartha Mitra, James Lennon, Sam Cheshier, Karen Elizabeth Lee, Jordan Roselli, Stephen Ahn, Grace Hagiwara, Mike Alvarez, Pieter Both, Jami Wang, Ben W Dulken, Maddalena Adorno, Vincent M. Alford, and Aisling Chaney. Flow cytometry analysis for this project was done on instruments in the Stanford Shared FACS Facility; BD FACSAriaII was purchased by NIH S10 shared instrumentation grant 1S10RR02933801. We thank the Stanford Neuroscience Microscopy Service, supported by NIH NS069375.

## Author contributions

FR, EC, BNdR, MMD and MFC conceptualized the aims of the project. Formal analysis, methodology, project administration, validation, visualization and writing the original draft was performed by FR, EC, and BNdR. The manuscript draft was edited by MFC, CH, MMD, EC, BNdR and FR. MFC contributed funding acquisition. Investigation was done by FR, EC, and BNdR. Resources were provided by MFC, SQ, AA, MMD and CH. CH supervised all behavioral testing, which was performed by BC. Real time experiments measuring reactive astrogliosis markers were performed by NG. VA and JA assisted with staining of pSMAD 1/5/8 and amyloid. RCJ and SK sequenced single-cell libraries and aligned reads. KP established human fetal neural stem cell lines for use in human AD model experiments. DQ performed breeding and oversaw mouse colonies.

## MATERIALS AND METHODS

### Statistical Analyses

In all the graphs, bars show average as central values and ± S.D. as error bars, unless otherwise specified. *P* values were calculated using ANOVA in analyses with 3 or mor groups. Two-tailed *t*-tests were used in analyses comparing 2 groups, unless otherwise specified. For limiting dilution analyses, ELDA software was used to test inequality between multiple groups. Expected frequencies are reported, as well as the 95% confidence intervals (lower and upper values are indicated). \**P*<0.05, \*\**P*<0.01, \*\*\**P* <0.001.

### Brain multianalyte analysis

The different brain regions were lysed using cell lysis buffer (Cell signaling #9803) with PMSF (Cell signaling #8553) and complete mini EDTA free protease inhibitor followed by mechanical homogenation by Tissue Ruptor (Qiagen). The samples were centrifuged at 13000 rpm for 15 mins and protein concentration calculated by BCA. Normalized samples were analyzed by the Stanford Human Immune Monitoring Center using a Luminex mouse 38-plex analyte platform that screens 38 secreted proteins using a multiplex fluorescent immunoassay. Brain homogenates were run in duplicate (three biological replicates were analyzed). The Luminex data (mean RFI) was generated by taking the raw fluorescence intensities of each sample and dividing by a control sample (one of the WT samples), then taking the average of the triplicated samples for each genotype.

### Flow Cytometry

For single-cell RNA-sequencing, the subventricular zone of 4 mice from each genotype was micro-dissected and tissue digested using Liberase DH (Roche) and DNAse I (250U/ml) at 37°C for 20 minutes followed by trituration. Digested tissue was washed in ice-cold HBSS without calcium and magnesium, filtered through a 40-μm filter, and then stained with the following antibodies for 30 minutes: PacBlue-CD31 (Biolegend), PacBlue-CD45 (Biolegend), PacBlue-Ter119 (Biolegend), and FITC-CD24 (Biolegend). Sytox Blue was used for cell death exclusion and samples were sorted into 384 well plates prepared with lysis buffer using the Sony Sorter.

### Lysis Plate Preparation

Lysis plates were created by dispensing 0.4 μl lysis buffer (0.5U Recombinant RNase Inhibitor (Takara Bio, 2313B), 0.0625% Triton™ X-100 (Sigma, 93443-100ML), 3.125 mM dNTP mix (Thermo Fisher, R0193), 3.125 μM Oligo-dT30VN (IDT, 5’AGCAGTGGTATCAACGCAGAGTACT30VN-3’) and 1:600,000 ERCC RNA spike-in mix (Thermo Fisher, 4456740)) into 384-well hard-shell PCR plates (Biorad HSP3901) using a Tempest or Mantis liquid handler (Formulatrix).

### cDNA Synthesis and Library Preparation

cDNA synthesis was performed using the Smart-seq2 protocol [1,2]. Illumina sequencing libraries were prepared according to the protocol in the Nextera XT Library Sample Preparation kit (Illumina, FC-131-1096). Each well was mixed with 0.8 μl Nextera tagmentation DNA buffer (Illumina) and 0.4 μl Tn5 enzyme (Illumina), then incubated at 55°C for 10 min. The reaction was stopped by adding 0.4 μl “Neutralize Tagment Buffer” (Illumina) and spinning at room temperature in a centrifuge at 3220xg for 5 min. Indexing PCR reactions were performed by adding 0.4 μl of 5 μM i5 indexing primer, 0.4 μl of 5 μM i7 indexing primer, and 1.2 μl of Nextera NPM mix (Illumina). PCR amplification was carried out on a ProFlex 2×384 thermal cycler using the following program: 1. 72°C for 3 minutes, 2. 95°C for 30 seconds, 3. 12 cycles of 95°C for 10 seconds, 55°C for 30 seconds, and 72°C for 1 minute, and 4. 72°C for 5 minutes.

### Library pooling, quality control, and sequencing

Following library preparation, wells of each library plate were pooled using a Mosquito liquid handler (TTP Labtech). Pooling was followed by two purifications using 0.7x AMPure beads (Fisher, A63881). Library quality was assessed using capillary electrophoresis on a Fragment Analyzer (AATI), and libraries were quantified by qPCR (Kapa Biosystems, KK4923) on a CFX96 Touch Real-Time PCR Detection System (Biorad). Plate pools were normalized to 2 nM and equal volumes from 10 or 20 plates were mixed together to make the sequencing sample pool. PhiX control library was spiked in at 0.2% before sequencing. Single-cell libraries were sequenced on the NovaSeq 6000 Sequencing System (Illumina) using 2 x 100bp paired-end reads and 2 x 8bp or 2 x 12bp index reads with a 300-cycle kit (Illumina 20012860).

### Data processing

Sequences were collected from the sequencer and de-multiplexed using bcl2fastq version 2.19.0.316. Reads were aligned using to the mm10plus genome using STAR version 2.5.2b with parameters TK. Gene counts were produced using HTSEQ version 0.6.1p1 with default parameters, except ‘stranded’ was set to ‘false’, and ‘mode’ was set to ‘intersection-nonempty’. As mentioned above, four biological replicates from each genotype and at each age were combined for the single-cell RNA-seq experiment (16 samples per age group). Basic filtering of cells and genes was conducted pre-analysis using the Seurat package in R (Butler, Hoffman, Smibert, Papalexi, & Satija, 2018). Briefly, genes that were not expressed in a minimum of 5 cells were filtered out, and cells had to have a minimum of 50,000 reads and a maximum of 3,000,000 reads. Similarly, cells with less than 500 or more than 5000 genes were filtered out. This left us with the following numbers of cells after filtering:

|  | 3-4 months | 1 year old |
| --- | --- | --- |
| WT | 1008 | 980 |
| Tg-SwDI | 642 | 1089 |
| Tg-SwDI/Usp16+/- | 712 | 729 |
| Usp16+/- | 651 | 731 |

### Gene Set Enrichment Analysis

Gene counts were log normalized and scaled before generating the .gct files. GSEA with the Hallmarks gene sets was run with standard parameters: 1000 permutations of type phenotype, with no collapsing to gene symbols, and weighted enrichment. Gene sets were considered significantly enriched if FDR<25%.

### Human neurosphere cultures

A human fetal neural stem cell line from University of California Irvine was developed from fetal neural tissue at 18 week gestational age enriched for CD133+ cells. The use of neural progenitor cells as non-hESC stem cells in this study is compliant to Stanford Stem Cell Research Oversight (SCRO) Protocol 194 pre-approved by the Internal Review Board (IRB)/SCRO of the Stanford Research Compliance Office (RCO). Informed consent was obtained, and standard material transfer agreement signed. Cells were grown in nonadherent ultra-low attachment well plates in X-VIVO 15 media (LONZA) supplemented with LIF (10 ng/ml), N2 Supplement, N-acetylcysteine (63 ug/ml), Heparin (2 ug/ml), EGF (20 ng/ml), and FGF (20 ng/ml).

For limiting dilution analysis, cells were directly plated into 96-well ultra-low adherent plates (Corning Costar) in limiting dilutions down to one cell per well. Each plating dose was done in replicates of up to 12 wells in each experiment, and the number of wells with neurospheres was counted after 10 days. Experiment was repeated 3 times.

### Lentivirus Production

cDNA for mutant APP harboring the Swedish and Indiana mutations was cloned into a pHIV-Zsgreen backbone obtained from Addgene. Lipofectamine 2000 was used to transduce the construct (either pHIV-Zsgreen+mutant APP or pHIV-Zsgreen alone) into H293T cells and media was collected after 48 hours. Virus was ultra-centrifuged and resuspended in PBS then titered before infecting human fetal neurospheres.

### Colony Counts

Human neurospheres were dissociated into single cells and infected with either a lentiviral construct containing pHIV-Zsgreen+mutant APP or pHIV-Zsgreen alone and allowed to grow for a week. Thereafter, cells were again dissociated and seeded at 5,000 cells/well in a 24-well plate in triplicate. Cells were fed every day with 20x media containing the appropriate amount of LDN-19389 (Selleckchem S2618). Colonies were counted after 7 days.

### Immunofluorescence

Neurospheres were cytospun onto slides and fixed in ice-cold methanol for 5 minutes. Slides were rinsed 3 times in phosphate-buffered saline (PBS) at room temperature, followed by blocking in 3% BSA in PBS for 1 hour at room temperature. Rabbit antibody to pSMAD 1/5/8 (1:100; CST 9516) and mouse antibody to beta-amyloid (1:100; Invitrogen 13-200) were diluted in the same 3% blocking buffer and incubated overnight at 4°C. The following day, sections were rinsed three times in 1X PBS and incubated in secondary antibody solution Cy-3 donkey anti-rabbit (1:500; Jackson ImmunoResearch) or Cy-3 donkey anti-mouse (1:500; Jackson ImmunoResearch) and 4’,6-diamidino-2-phenylindole (DAPI) (1:10,000) in 3% blocking solution at room temperature for 2 hours. Slides were then washed 3 times at room temperature in 1X PBS and mounted.

### Mice

Tg-SwDI mice (background C57Bl/6) were purchased from Jackson Laboratories. These mice were made hemizygous for experiments after breeding with *Cdkn2a^-/-^* (C57Bl6 background) or *Usp16^+/-^* mice (back-crossed to B6EiC3). *Usp16^+/-^* mice were originally ordered from Mutant Mouse Regional Resource Centers (MMRRC) and *Cdkn2a^-/-^*(B6.129-*Cdkn2a^tm1Rdp^*) were obtained from Mouse Models of Human Cancers Consortium (NCI-Frederick). Wild-type littermates were used as control mice. Mice were genotyped by traditional PCR according to animal’s provider. Mice were housed in accordance with the guidelines of Institutional Animal Care Use Committee. All animal procedures and behavioral studies involved in this manuscript are compliant to Stanford Administrative Panel on Laboratory Animal Care (APLAC) Protocol 10868 pre-approved by the Stanford Institutional Animal Care and Use Committee (IACUC).

### Mouse neurosphere cultures

To produce neurospheres, mice were euthanized by CO2, decapitated and the brain immediately removed. The subventricular zone was micro-dissected and stored in ice-cold PBS for further processing. The tissue was digested using Liberase DH (Roche) and DNAse I (250U/ml) at 37°C for 20 minutes followed by trituration. Digested tissue was washed in ice-cold HBSS without calcium and magnesium, filtered through a 40-μm filter and immediately put into neurosphere growth media that is, Neurobasal-A (Invitrogen) supplemented with Glutamax (Life Technologies), 2% B27-A (Invitrogen), mouse recombinant epidermal growth factor (EGF; 20 ng/ml) and basic fibroblast growth factor (bFGF; 20 ng/ml) (Shenandoah Biotechnology).

For limiting dilution analysis, cells were directly plated into 96-well ultra-low adherent plates (Corning Costar) in limiting dilutions down to one cell per well. Each plating dose was done in replicate of up to 12 wells in each experiment, and the number of wells with neurospheres was counted after 10 days. For passaging, neurospheres were dissociated and re-plated at a density of 10 cells/uL.

### RNA expression analyses (mouse)

For real-time analyses, cells were collected in Trizol (Invitrogen), and RNA was extracted following the manufacturer’s protocol. Complementary DNA was obtained using Superscript III First Strand Synthesis (Invitrogen). Real-time reactions were assembled using Taqman probes (Applied Biosystems) in accordance with the manufacturer’s directions. Expression data were normalized by the expression of housekeeping gene ActB (Mm00607939_s1). Probes used in this study: Cdkn2a (Mm_00494449), Bmi1 (Mm03053308_g1), IL1ß (Mm01336189_m1), IL6 (Mm99999064_m1), TNF (Mm00443258_m1) and COX2 (Mm03294838_g1).

### Immunohistochemistry

All animals were anesthetized with avertin and transcardially perfused with 15 ml phosphate-buffered saline (PBS). Brains were postfixed in 4% paraformaldehyde (PFA) overnight at 4°C before cryoprotection in 30% sucrose. Brains were embedded in optimum cutting temperature (Tissue-Tek) and coronally sectioned at 40 μm using a sliding microtome (Leica, HM450). For immunohistochemistry, sections were stained using the Click-iT EdU cell proliferation kit and protocol (Invitrogen) to expose EdU labeling followed by incubation in blocking solution [3% normal donkey serum, 0.3% Triton X-100 in phosphate-buffered saline (PBS)] at room temperature for 1 hour. Goat antibody to Sox2 (anti-Sox2) (1:50; R&D Systems AF2018) and rabbit anti-GFAP (1:500; Stem Cell Technologies 60128) were diluted in 1% blocking solution (normal donkey serum in 0.3% Triton X-100 in PBS) and incubated overnight at 4°C. Secondary-only stains were performed as negative controls. The following day, sections were rinsed three times in 1X PBS and incubated in secondary antibody solution (1:500) and 4’,6-diamidino-2-phenylindole (DAPI) (1:10,000) in 1% blocking solution at 4°C for 4 hours. The following secondary antibodies were used: Alexa 594 donkey anti-rabbit (Jackson ImmunoResearch), Alexa 647 donkey anti-goat (Jackson ImmunoResearch). The next day, sections were rinsed three times in PBS and mounted with ProLong Gold Antifade (Cell Signaling) mounting medium. For senile plaques, sections were incubated for 8 min in aqueous 1% Thioflavin S (Sigma) at room temperature, washed in ethanol and mounted. Total plaque area from images taken of 6 sections were analyzed from each mouse with n=3 mice in each group.

### Confocal Imaging and Quantification

All cell counting was performed by experimenters blinded to the experimental conditions using a Zeiss LSM700 scanning confocal microscope (Carl Zeiss). For EdU stereology, all EdU-labeled cells in every 6th coronal section of the SVZ were counted by blinded experimenters at 40× magnification. The total number of EdU-labeled cells co-labeled with Sox2 and GFAP per SVZ was determined by multiplying the number of EdU^+^GFAP^+^Sox2^+^ cells by 6. Cells were considered triple-labeled when they colocalized within the same plane.

### Behavioral testing

*Novel object recognition:* One behavioral test used in this study for assessing long term memory was novel object recognition (NOR) ^67^ carried out in arenas (50cm x 50cm x 50cm) resting on an infra-red emitting base. Behavior was recorded by an infrared-sensitive camera placed 2.5m above the arena. Data were stored and analyzed using Videotrack software from ViewPoint Life Sciences, Inc. (Montreal, Canada) allowing the tracking of body trajectory/speed and the detection of the nose position. On the day before NOR training, the mouse was habituated to the apparatus by freely exploring the open arena. NOR is based on the preference of mice for a novel object versus a familiar object when allowed to explore freely. For NOR training, two identical objects were placed into the arena and the animals were allowed to explore for 10 minutes. Testing occurred 24 hours later in the same arena but one of the familiar objects used during training was replaced by a novel object of similar dimensions, and the animal was allowed to explore freely for 7 min. The objects and the arena were cleaned with 10% ethanol between trials. Exploration of the objects was defined by the time spent with the nose in a 2.5cm zone around the objects. The preference index (P.I.) was calculated as the ratio of the time spent exploring the novel object over the total time spent exploring the two objects. The P.I. was calculated for each animal and averaged among the groups of mice by genotype. The P.I. should not be significantly different from 50% in the training session, but is significantly different if novelty is detected.

*Barnes Maze:* Another test of long-term memory that is indicative of spatial memory is the Barnes Maze similar to that described by Attar et al. (Attar et al., 2013). The Barnes maze is a 20-hole circular platform measuring 48” in diameter with holes cut 1.75” in diameter and 1” from the edge. The platform is elevated 100 cm above the floor, and is located in the center of a room with many extra-maze and intra-maze visual cues. This task takes advantage of the natural preference of rodents for a dark environment. Motivated to escape the bright lights and the open-space of the platform, rodents search for an escape hole that leads to a dark box beneath the maze and with training they learn to use distal visual cues to determine the spatial location of the escape hole. A habituation day was followed by training over 2 days and a test day separated by 24hrs. Two downward-facing 150-watt incandescent light bulbs mounted overhead served as an aversive stimulus. Mice completed three phases of testing: habituation, training, and the probe test.

For habituation, mice were placed within the start cylinder in the middle of the maze to ensure random orientation for 15 sec. The overhead lights were then turned on and mice were given 3 min to independently enter through the target hole into the escape cage. If a mouse did not enter the escape box freely, the experimenter coaxed the mouse to enter the escape box by touching the mouse’s tail.

For training, a mouse was placed in the middle of the maze in random orientation for 15 sec. The overhead lights were turned on, and the tracking software was activated. The mouse was allowed up to 3 minutes to explore the maze and enter the escape hole. If it failed to enter within 3 minutes, it was gently guided to the escape hole using the start cylinder and allowed to enter the escape cage independently.

On the test/probe day, 24 hours after the last training day, the experiment was set up as described on training days, except the target hole was covered. The percent time in the correct zone and average proximity to the correct escape hole are more sensitive measures of memory than percentage visits to the correct hole. Therefore, during the probe phase, measures of time spent per quadrant and holes searched per quadrant were recorded. For these analyses, the maze was divided into quadrants consisting of 5 holes with the target hole in the center of the target quadrant. On day 4, latency (seconds) and path length (meters) to reach the target hole were measured. Number of pokes in each hole were calculated, time spent per quadrant and holes searched per quadrant were recorded and paired t-tests were used to compare the percentage of time spent between quadrants.

## REFERENCES AND NOTES

Abdouh, M., Chatoo, W., El Hajjar, J., David, J., Ferreira, J., & Bernier, G. (2012). Bmi1 is down-regulated in the aging brain and displays antioxidant and protective activities in neurons. PLoS One, 7(2), e31870. DOI: 10.1371/journal.pone.0031870.

Adorno, M., Sikandar, S., Mitra, S. S., Kuo, A., Nicolis Di Robilant, B., Haro-Acosta, V., . . . Clarke, M. F. (2013). Usp16 contributes to somatic stem-cell defects in Down’s syndrome. Nature, 501(7467), 380–384. DOI: 10.1038/nature12530.

Aisen, P. S. (2008). Tarenflurbil: a shot on goal. Lancet Neurol, 7(6), 468–469. DOI: 10.1016/S1474-4422(08)70091-7.

Akiyama, H., Barger, S., Barnum, S., Bradt, B., Bauer, J., Cole, G. M., . . . Wyss-Coray, T. (2000). Inflammation and Alzheimer’s disease. Neurobiol Aging, 21(3), 383–421. DOI: 10.1016/s0197-4580(00)00124-x.

Alipour, M., Nabavi, S. M., Arab, L., Vosough, M., Pakdaman, H., Ehsani, E., & Shahpasand, K. (2019). Stem cell therapy in Alzheimer’s disease: possible benefits and limiting drawbacks. Mol Biol Rep, 46(1), 1425–1446. DOI: 10.1007/s11033-018-4499-7.

Amaya-Montoya, M., Perez-Londono, A., Guatibonza-Garcia, V., Vargas-Villanueva, A., & Mendivil, C. O. (2020). Cellular Senescence as a Therapeutic Target for Age-Related Diseases: A Review. Adv Ther, 37(4), 1407–1424. DOI: 10.1007/s12325-020-01287-0.

Attar, A., Liu, T., Chan, W. T., Hayes, J., Nejad, M., Lei, K., & Bitan, G. (2013). A shortened Barnes maze protocol reveals memory deficits at 4-months of age in the triple-transgenic mouse model of Alzheimer’s disease. PLoS One, 8(11), e80355. DOI: 10.1371/journal.pone.0080355.

Baker, D. J., Wijshake, T., Tchkonia, T., LeBrasseur, N. K., Childs, B. G., van de Sluis, B., . . . van Deursen, J. M. (2011). Clearance of p16Ink4a-positive senescent cells delays ageing-associated disorders. Nature, 479(7372), 232–236. DOI: 10.1038/nature10600.

Bruggeman, S. W., Valk-Lingbeek, M. E., van der Stoop, P. P., Jacobs, J. J., Kieboom, K., Tanger, E., . . . van Lohuizen, M. (2005). Ink4a and Arf differentially affect cell proliferation and neural stem cell self-renewal in Bmi1-deficient mice. Genes Dev, 19(12), 1438–1443. DOI: 10.1101/gad.1299305.

Bussian, T. J., Aziz, A., Meyer, C. F., Swenson, B. L., van Deursen, J. M., & Baker, D. J. (2018). Clearance of senescent glial cells prevents tau-dependent pathology and cognitive decline. Nature, 562(7728), 578–582. DOI: 10.1038/s41586-018-0543-y.

Butler, A., Hoffman, P., Smibert, P., Papalexi, E., & Satija, R. (2018). Integrating single-cell transcriptomic data across different conditions, technologies, and species. Nat Biotechnol, 36(5), 411–420. DOI: 10.1038/nbt.4096.

Castellani, R. J., Rolston, R. K., & Smith, M. A. (2010). Alzheimer disease. Dis Mon, 56(9), 484–546. DOI: 10.1016/j.disamonth.2010.06.001.

Chang, J., Dettman, R. W., & Dizon, M. L. V. (2018). Bone morphogenetic protein signaling: a promising target for white matter protection in perinatal brain injury. Neural Regen Res, 13(7), 1183–1184. DOI: 10.4103/1673-5374.235025.

Chehrehasa, F., Meedeniya, A. C., Dwyer, P., Abrahamsen, G., & Mackay-Sim, A. (2009). EdU, a new thymidine analogue for labelling proliferating cells in the nervous system. J Neurosci Methods, 177(1), 122–130. DOI: 10.1016/j.jneumeth.2008.10.006.

Davis, J., Xu, F., Deane, R., Romanov, G., Previti, M. L., Zeigler, K., . . . Van Nostrand, W. E. (2004). Early-onset and robust cerebral microvascular accumulation of amyloid beta-protein in transgenic mice expressing low levels of a vasculotropic Dutch/Iowa mutant form of amyloid beta-protein precursor. J Biol Chem, 279(19), 20296–20306. DOI: 10.1074/jbc.M312946200.

Doody, R. S., Raman, R., Farlow, M., Iwatsubo, T., Vellas, B., Joffe, S., . . . Semagacestat Study, G. (2013). A phase 3 trial of semagacestat for treatment of Alzheimer’s disease. N Engl J Med, 369(4), 341–350. DOI: 10.1056/NEJMoa1210951.

Ennaceur, A., & Delacour, J. (1988). A new one-trial test for neurobiological studies of memory in rats. 1: Behavioral data. Behav Brain Res, 31(1), 47–59. DOI: 10.1016/0166-4328(88)90157-x.

Essers, M. A., Offner, S., Blanco-Bose, W. E., Waibler, Z., Kalinke, U., Duchosal, M. A., & Trumpp, A. (2009). IFNalpha activates dormant haematopoietic stem cells in vivo. Nature, 458(7240), 904–908. DOI: 10.1038/nature07815.

Flamier, A., El Hajjar, J., Adjaye, J., Fernandes, K. J., Abdouh, M., & Bernier, G. (2018). Modeling Late-Onset Sporadic Alzheimer’s Disease through BMI1 Deficiency. Cell Rep, 23(9), 2653–2666. DOI: 10.1016/j.celrep.2018.04.097.

Frost, G. R., & Li, Y. M. (2017). The role of astrocytes in amyloid production and Alzheimer’s disease. Open Biol, 7(12). DOI: 10.1098/rsob.170228.

Gargiulo, G., Cesaroni, M., Serresi, M., de Vries, N., Hulsman, D., Bruggeman, S. W., . . . van Lohuizen, M. (2013). In vivo RNAi screen for BMI1 targets identifies TGF-beta/BMP-ER stress pathways as key regulators of neural-and malignant glioma-stem cell homeostasis. Cancer Cell, 23(5), 660–676. DOI: 10.1016/j.ccr.2013.03.030.

Gjoneska, E., Pfenning, A. R., Mathys, H., Quon, G., Kundaje, A., Tsai, L. H., & Kellis, M. (2015). Conserved epigenomic signals in mice and humans reveal immune basis of Alzheimer’s disease. Nature, 518(7539), 365–369. DOI: 10.1038/nature14252.

Glass, C. K., Saijo, K., Winner, B., Marchetto, M. C., & Gage, F. H. (2010). Mechanisms underlying inflammation in neurodegeneration. Cell, 140(6), 918–934. DOI: 10.1016/j.cell.2010.02.016.

Green, R. C., Schneider, L. S., Amato, D. A., Beelen, A. P., Wilcock, G., Swabb, E. A., . . . Tarenflurbil Phase 3 Study, G. (2009). Effect of tarenflurbil on cognitive decline and activities of daily living in patients with mild Alzheimer disease: a randomized controlled trial. JAMA, 302(23), 2557–2564. DOI: 10.1001/jama.2009.1866.

Group, A. D. C., Bentham, P., Gray, R., Sellwood, E., Hills, R., Crome, P., & Raftery, J. (2008). Aspirin in Alzheimer’s disease (AD2000): a randomised open-label trial. Lancet Neurol, 7(1), 41–49. DOI: 10.1016/S1474-4422(07)70293-4.

Haughey, N. J., Liu, D., Nath, A., Borchard, A. C., & Mattson, M. P. (2002). Disruption of neurogenesis in the subventricular zone of adult mice, and in human cortical neuronal precursor cells in culture, by amyloid beta-peptide: implications for the pathogenesis of Alzheimer’s disease. Neuromolecular Med, 1(2), 125–135. DOI: 10.1385/NMM:1:2:125.

Hebert, L. E., Weuve, J., Scherr, P. A., & Evans, D. A. (2013). Alzheimer disease in the United States (2010-2050) estimated using the 2010 census. Neurology, 80(19), 1778–1783. DOI: 10.1212/WNL.0b013e31828726f5.

Hu, Y., & Smyth, G. K. (2009). ELDA: extreme limiting dilution analysis for comparing depleted and enriched populations in stem cell and other assays. J Immunol Methods, 347(1-2), 70–78. DOI: 10.1016/j.jim.2009.06.008.

Huang, Y., & Mucke, L. (2012). Alzheimer mechanisms and therapeutic strategies. Cell, 148(6), 1204–1222. DOI: 10.1016/j.cell.2012.02.040.

Hussussian, C. J., Struewing, J. P., Goldstein, A. M., Higgins, P. A., Ally, D. S., Sheahan, M. D., . . . Dracopoli, N. C. (1994). Germline p16 mutations in familial melanoma. Nat Genet, 8(1), 15–21. DOI: 10.1038/ng0994-15.

Imbimbo, B. P., Solfrizzi, V., & Panza, F. (2010). Are NSAIDs useful to treat Alzheimer’s disease or mild cognitive impairment? Front Aging Neurosci, 2. DOI: 10.3389/fnagi.2010.00019.

Joo, H. Y., Zhai, L., Yang, C., Nie, S., Erdjument-Bromage, H., Tempst, P., . . . Wang, H. (2007). Regulation of cell cycle progression and gene expression by H2A deubiquitination. Nature, 449(7165), 1068–1072. DOI: 10.1038/nature06256.

Krishnamurthy, J., Ramsey, M. R., Ligon, K. L., Torrice, C., Koh, A., Bonner-Weir, S., & Sharpless, N. E. (2006). p16INK4a induces an age-dependent decline in islet regenerative potential. Nature, 443(7110), 453–457. DOI: 10.1038/nature05092.

Kwak, Y. D., Hendrix, B. J., & Sugaya, K. (2014). Secreted type of amyloid precursor protein induces glial differentiation by stimulating the BMP/Smad signaling pathway. Biochem Biophys Res Commun, 447(3), 394–399. DOI: 10.1016/j.bbrc.2014.03.139.

Leeman, D. S., Hebestreit, K., Ruetz, T., Webb, A. E., McKay, A., Pollina, E. A., . . . Brunet, A. (2018). Lysosome activation clears aggregates and enhances quiescent neural stem cell activation during aging. Science, 359(6381), 1277–1283. DOI: 10.1126/science.aag3048.

Lopez-Toledano, M. A., & Shelanski, M. L. (2004). Neurogenic effect of beta-amyloid peptide in the development of neural stem cells. J Neurosci, 24(23), 5439–5444. DOI: 10.1523/JNEUROSCI.0974-04.2004.

Meyers, E. A., Gobeske, K. T., Bond, A. M., Jarrett, J. C., Peng, C. Y., & Kessler, J. A. (2016). Increased bone morphogenetic protein signaling contributes to age-related declines in neurogenesis and cognition. Neurobiol Aging, 38, 164–175. DOI: 10.1016/j.neurobiolaging.2015.10.035.

Miao, J., Xu, F., Davis, J., Otte-Holler, I., Verbeek, M. M., & Van Nostrand, W. E. (2005). Cerebral microvascular amyloid beta protein deposition induces vascular degeneration and neuroinflammation in transgenic mice expressing human vasculotropic mutant amyloid beta precursor protein. Am J Pathol, 167(2), 505–515. DOI: 10.1016/s0002-9440(10)62993-8.

Molofsky, A. V., Pardal, R., Iwashita, T., Park, I. K., Clarke, M. F., & Morrison, S. J. (2003). Bmi-1 dependence distinguishes neural stem cell self-renewal from progenitor proliferation. Nature, 425(6961), 962–967. DOI: 10.1038/nature02060.

Molofsky, A. V., Slutsky, S. G., Joseph, N. M., He, S., Pardal, R., Krishnamurthy, J., . . . Morrison, S. J. (2006). Increasing p16INK4a expression decreases forebrain progenitors and neurogenesis during ageing. Nature, 443(7110), 448–452. DOI: 10.1038/nature05091.

Mootha, V. K., Lindgren, C. M., Eriksson, K. F., Subramanian, A., Sihag, S., Lehar, J., . . . Groop, L. C. (2003). PGC-1alpha-responsive genes involved in oxidative phosphorylation are coordinately downregulated in human diabetes. Nat Genet, 34(3), 267–273. DOI: 10.1038/ng1180.

Mu, Y., & Gage, F. H. (2011). Adult hippocampal neurogenesis and its role in Alzheimer’s disease. Mol Neurodegener, 6, 85. DOI: 10.1186/1750-1326-6-85.

Neuropathology Group. Medical Research Council Cognitive, F., & Aging, S. (2001). Pathological correlates of late-onset dementia in a multicentre, community-based population in England and Wales. Neuropathology Group of the Medical Research Council Cognitive Function and Ageing Study (MRC CFAS). Lancet, 357(9251), 169–175. DOI: 10.1016/s0140-6736(00)03589-3.

Osborn, L. M., Kamphuis, W., Wadman, W. J., & Hol, E. M. (2016). Astrogliosis: An integral player in the pathogenesis of Alzheimer’s disease. Prog Neurobiol, 144, 121–141. DOI: 10.1016/j.pneurobio.2016.01.001.

Pastrana, E., Silva-Vargas, V., & Doetsch, F. (2011). Eyes wide open: a critical review of sphere-formation as an assay for stem cells. Cell Stem Cell, 8(5), 486–498. DOI: 10.1016/j.stem.2011.04.007.

Rodriguez, J. J., Jones, V. C., & Verkhratsky, A. (2009). Impaired cell proliferation in the subventricular zone in an Alzheimer’s disease model. Neuroreport, 20(10), 907–912. DOI: 10.1097/WNR.0b013e32832be77d.

Rodriguez, J. J., & Verkhratsky, A. (2011). Neurogenesis in Alzheimer’s disease. J Anat, 219(1), 78–89. DOI: 10.1111/j.1469-7580.2011.01343.x.

Sakamoto, M., Ieki, N., Miyoshi, G., Mochimaru, D., Miyachi, H., Imura, T., . . . Imayoshi, I. (2014). Continuous postnatal neurogenesis contributes to formation of the olfactory bulb neural circuits and flexible olfactory associative learning. J Neurosci, 34(17), 5788–5799. DOI: 10.1523/JNEUROSCI.0674-14.2014.

Salloway, S., Sperling, R., Fox, N. C., Blennow, K., Klunk, W., Raskind, M., . . . Clinical Trial, I. (2014). Two phase 3 trials of bapineuzumab in mild-to-moderate Alzheimer’s disease. N Engl J Med, 370(4), 322–333. DOI: 10.1056/NEJMoa1304839.

Scheeren, F. A., Kuo, A. H., van Weele, L. J., Cai, S., Glykofridis, I., Sikandar, S. S., . . . Clarke, M. F. (2014). A cell-intrinsic role for TLR2-MYD88 in intestinal and breast epithelia and oncogenesis. Nat Cell Biol, 16(12), 1238–1248. DOI: 10.1038/ncb3058.

Selkoe, D. J. (2019). Alzheimer disease and aducanumab: adjusting our approach. Nat Rev Neurol, 15(7), 365–366. DOI: 10.1038/s41582-019-0205-1.

Subramanian, A., Tamayo, P., Mootha, V. K., Mukherjee, S., Ebert, B. L., Gillette, M. A., . . . Mesirov, J. P. (2005). Gene set enrichment analysis: a knowledge-based approach for interpreting genome-wide expression profiles. Proc Natl Acad Sci U S A, 102(43), 15545–15550. DOI: 10.1073/pnas.0506580102.

Thal, D. R., Ghebremedhin, E., Orantes, M., & Wiestler, O. D. (2003). Vascular pathology in Alzheimer disease: correlation of cerebral amyloid angiopathy and arteriosclerosis/lipohyalinosis with cognitive decline. J Neuropathol Exp Neurol, 62(12), 1287–1301. DOI: 10.1093/jnen/62.12.1287.

Wilcock, D. M., Lewis, M. R., Van Nostrand, W. E., Davis, J., Previti, M. L., Gharkholonarehe, N., . . . Colton, C. A. (2008). Progression of amyloid pathology to Alzheimer’s disease pathology in an amyloid precursor protein transgenic mouse model by removal of nitric oxide synthase 2. J Neurosci, 28(7), 1537–1545. DOI: 10.1523/JNEUROSCI.5066-07.2008.

Winner, B., Kohl, Z., & Gage, F. H. (2011). Neurodegenerative disease and adult neurogenesis. Eur J Neurosci, 33(6), 1139–1151. DOI: 10.1111/j.1460-9568.2011.07613.x.

Yousef, H., Morgenthaler, A., Schlesinger, C., Bugaj, L., Conboy, I. M., & Schaffer, D. V. (2015). Age-Associated Increase in BMP Signaling Inhibits Hippocampal Neurogenesis. Stem Cells, 33(5), 1577–1588. DOI: 10.1002/stem.1943.

Yu, P. B., Deng, D. Y., Lai, C. S., Hong, C. C., Cuny, G. D., Bouxsein, M. L., . . . Bloch, K. D. (2008). BMP type I receptor inhibition reduces heterotopic [corrected] ossification. Nat Med, 14(12), 1363–1369. DOI: 10.1038/nm.1888.

Zencak, D., Lingbeek, M., Kostic, C., Tekaya, M., Tanger, E., Hornfeld, D., . . . Arsenijevic, Y. (2005). Bmi1 loss produces an increase in astroglial cells and a decrease in neural stem cell population and proliferation. J Neurosci, 25(24), 5774–5783. DOI: 10.1523/JNEUROSCI.3452-04.2005.

Zhu, Y., Tchkonia, T., Pirtskhalava, T., Gower, A. C., Ding, H., Giorgadze, N., . . . Kirkland, J. L. (2015). The Achilles’ heel of senescent cells: from transcriptome to senolytic drugs. Aging Cell, 14(4), 644–658. DOI: 10.1111/acel.12344.

